# Modeling the Depth of Cellular Dormancy from RNA-sequencing data

**DOI:** 10.1101/2023.12.11.571187

**Authors:** Michelle Yuchen Wei, Guang Yao

## Abstract

High-throughput transcriptome RNA sequencing is a powerful tool for understanding dynamic biological processes. Here, we present a computational framework, implemented in an R package QDSWorkflow, to characterize heterogeneous cellular dormancy depth using RNA-sequencing data from bulk samples and single cells.

## 1 Introduction

The vast majority of cells in the human body are not actively dividing but dormant [1]. Among these dormant cells, some are reversible (quiescent) and can reenter the cell cycle upon growth stimulation, whereas others are irreversibly arrested in senescence or terminal differentiation. Often referred to as G0, cellular dormancy states exhibit significant heterogeneity in "depth" [2, 3]. In quiescence, deeper quiescent cells, although fully reversible, require stronger growth stimulation and take a longer time to reenter the cell cycle than shallower quiescent cells. Examples of quiescence deepening have long been observed in cells under extended periods of serum starvation or contact inhibition in culture and with aging *in vivo* [4-10]. On the contrary, subpopulations of quiescent muscle, neural, and hematopoietic stem cells, upon tissue injury or in response to systemic circulating factors, move to shallower (primed or GAlert) quiescence that bear a closer resemblance to active cells [11-14]. Similarly, senescent cells also exhibit different "depths" with different cellular characteristics [15].

Our recent work suggests that quiescence deepening represents a transition trajectory from active proliferation to permanent senescence, and quiescence depth is regulated by cellular dimmer switch-like mechanisms [16-18]. Consistently, we found that a gene signature to predict quiescence depth score (QDS) in cultured fibroblasts can correctly predict a wide array of senescent and aging cell types in publicly available datasets [16]. To streamline the computational process to model the QDS of cellular dormancy states, we created a QDSWorkflow R package. Here, we present the methods for using QDSWorkflow to build an elastic net regression model and predict the relative dormancy depths of cells in growth, quiescence, and senescence from bulk and single-cell RNA-sequencing data.

## 2 Materials

### 2.1 Statistical modeling and machine learning

1. A computer with Windows, Mac, or Linux operating system.
2. R program (version 4.2 or higher, see note 1).
3. RStudio program (see note 1).
4. QDSWorkflow R package (see note 2).

### 2.2 RNA-seq datasets of cellular dormancy

1. Training data. An example dataset used in this chapter is a bulk RNA-seq dataset of rat embryonic fibroblasts (REF) under 2-16 days of serum starvation and displaying a continuously increased quiescence depth over this time window [16].
2. Test data. Example datasets used here are a) bulk RNA-seq of human HFF fibroblasts from 16-74 population doublings [19], b) bulk RNA-seq of growing, quiescent, senescent, and deep senescent human BJ fibroblasts [20], and c) scRNA-seq of quiescent and activated mouse neural stem cells (NSC) [21].

## 3 Methods

### 3.1 Computer setup

1. Download and install R (see note 1).
2. Download and install RStudio (see note 1).
3. Download and install the QDSWorkflow R package, which performs the modeling steps in the workflow shown in Fig. 1 (see note 2).

**Fig. 1.**
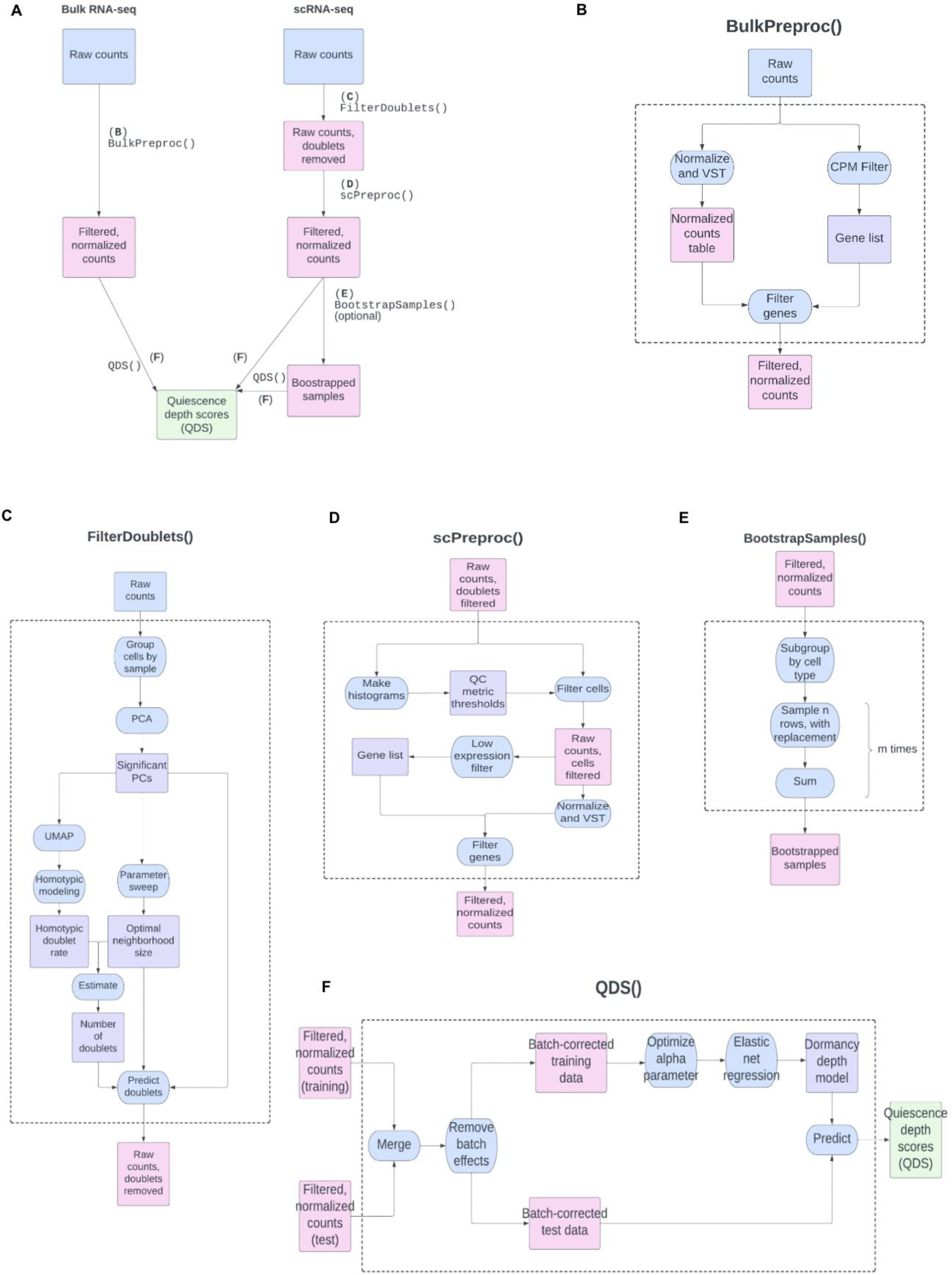
Flowchart to model cell dormancy depth using the QDSWorkflow package. A. Main workflow. Individual function modules (B-F) are labeled by function names. B-F. Function modules. Shown in the dashed box are the detailed steps of each function module. See Notes for details.

### 3.2 Data preprocessing for bulk RNA-seq datasets

1. Load the raw counts sequencing data into R (see note 3).
2. Normalize raw counts to eliminate systematic variations between sequencing samples (e.g., sequencing depth, library composition, etc.) and filter out lowly or non-expressed genes by running the BulkPreproc() function (Fig. 1B; see note 4).

### 3.3 Data preprocessing for single-cell RNA-seq (scRNA-seq) datasets

1. Load the raw counts sequencing data into R (see note 3).
2. Remove doublet reads (when multiple cells are sequenced as one cell) by running the FilterDoublets() function (Fig. 1C; see note 5).
3. Choose threshold values for cell reads QC metrics (total counts, gene number, mitochondrial ratio, etc.) based on QC value distribution histograms (see note 6).
4. Filter out low-quality cells using the chosen QC thresholds above, normalize the counts, and filter out lowly or non-expressed genes by running the scPreproc() function (Fig. 1D; see note 7).

### 3.4 Predict cell dormancy depth using a regression model

1. Load the training RNA-seq data into R (see note 8). This training dataset should be associated with different cell dormancy depths (e.g., dataset in 2.2.1) and preprocessed using the steps in 3.2.
2. Load the test data into R. Here, we use three example datasets (2.2.2), which are preprocessed using the steps in 3.2 (bulk RNA-seq data) or 3.3 (scRNA-seq).
3. Match the genes between the training and test datasets. When necessary, convert gene names and IDs using the convertSymbol() function (see note 9) and convert gene IDs between species using the convertSpecies() function (see note 10). For example, we convert the gene Ensembl IDs in the training data from rat to human and mouse, respectively, to match the human gene IDs in the test datasets 2.2.2a and 2.2.2b and mouse gene IDs in the test dataset 2.2.2c.
4. (Optional) For scRNA-seq test data that are typically sparse in gene expression values, make binned bootstrapped samples using the BootstrapSamples() function (Fig. 1E; see note 11). Binning cells in scRNA-seq often makes the trend in cell dormancy depth clearer.
5. Build a linear regression model using the training data and make predictions on the test data by running the QDS() function (Fig. 1F; see note 12). The QDS() function first merges the training and test datasets and uses the ComBat() function in the sva package [22] to remove batch effects between them. Then, it builds an elastic net regression model [23]on the training data by optimizing the cross-validation error (see note 13). For a given test set, a table of predicted quiescence depth scores (QDS) associated with individual samples is generated, with the QDS values indicating the relative dormancy depth difference across samples in the test dataset.
6. Visualize the predicted quiescence depth scores (QDS). To make boxplots of the predicted QDS values, use the make_boxplot() function if there is one sample group per condition (Fig. 2 & 3; see note 14) or the make_grouped_boxplot() function if there is more than one sample group per condition (Fig. 4; see note 15). For scRNA-seq data, density curves can be plotted using the make_grouped_hist() function to visualize the distribution of QDS values of individual cells within a population (Fig. 5; see note 16).

**Fig. 2.**
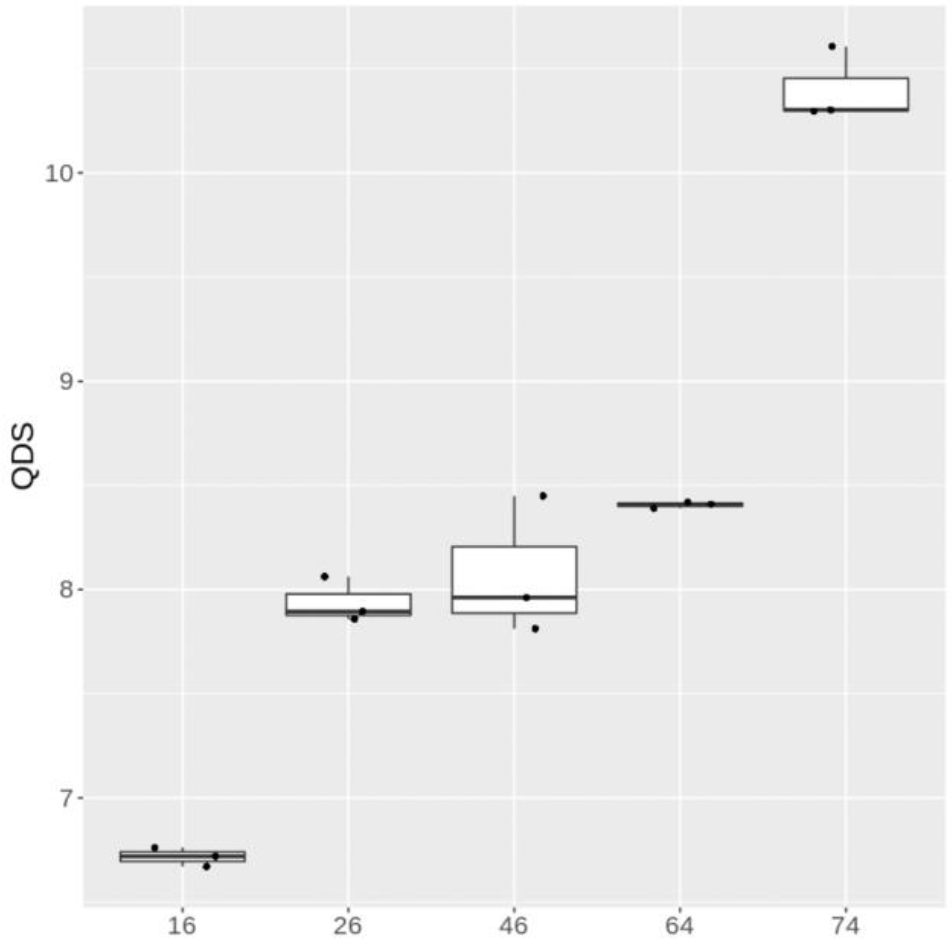
Predicted QDS of human HFF fibroblasts from 16 to 74 population doublings. The QDS values increase with population doublings, which reflects the decrease in proliferative activity and potential as cells approach replicative senescence. Each sample group contains triplicate samples at a population doubling [19].

**Fig. 3.**
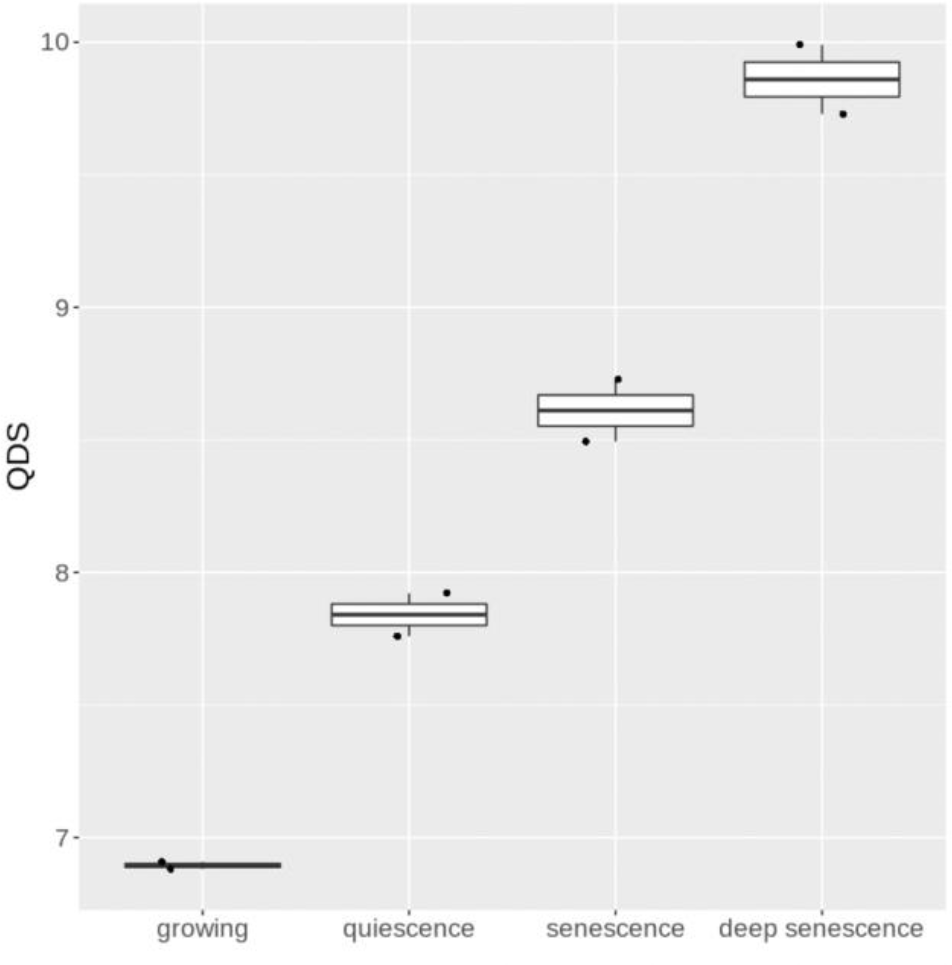
Predicted QDS of human BJ fibroblasts under growing, quiescent, senescent, and deep senescent conditions. Quiescence was induced by serum starvation. Senescence and deep senescence were induced under bleomycin treatment for 2 hours, then culturing for 12 and 28 days, respectively. Note that the QDS values keep increasing from growing to quiescence, to senescence, and then to deep senescence, reflecting continuously increased dormancy depth and decreased proliferation activity and potential. Each sample group contains duplicate samples at a cell state [20].

**Fig. 4.**
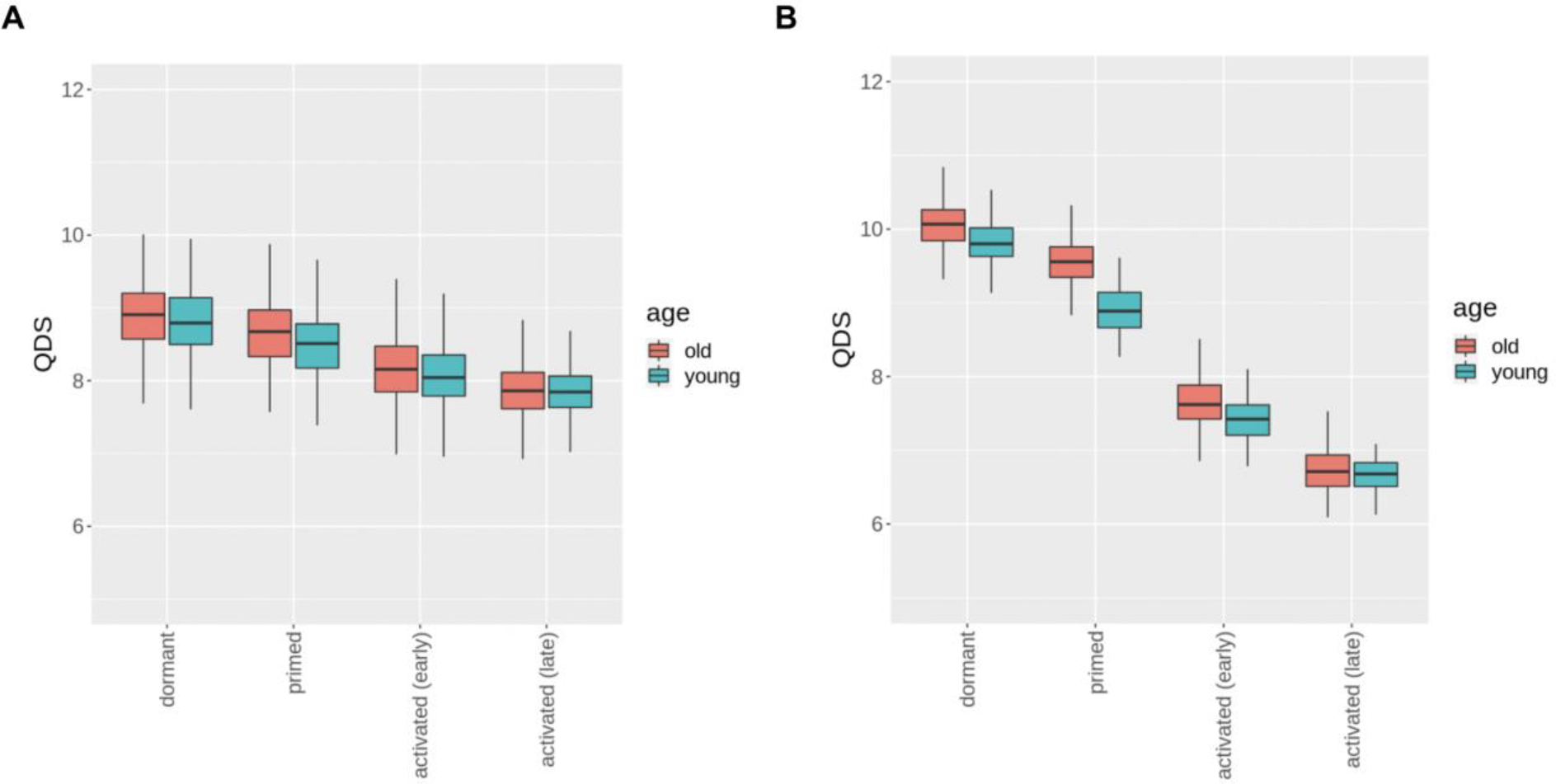
Predicted QDS of mouse neural stem cells from dormancy to activation. A. QDS values for individual cells in a population at the indicated condition. B. QDS values for bootstrapped samples (100 bootstrapped samples of 50 cells each were created for each condition). In both plots, the predicted QDS values are higher in old NSCs than in young NSCs, indicating the increased quiescence depth with aging. The QDS values decrease as cells gradually progress from dormancy to activation, reflecting the increase in proliferative potential and activity. The scRNA-seq dataset and the cell type labels are obtained from [21].

**Fig. 5.**
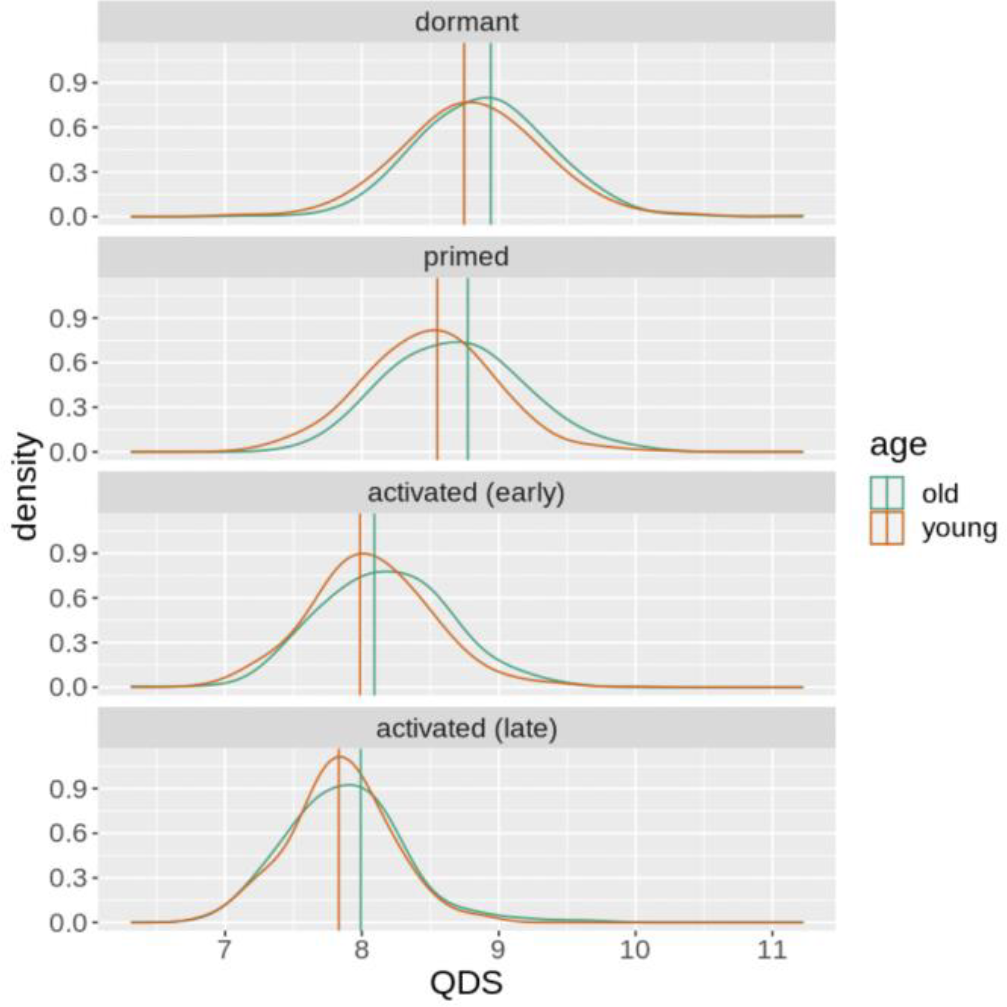
QDS density curves. Each curve represents the density distribution of QDS values within a cell population in the mouse NSC dataset [21]. The vertical lines show the mode of each curve. Old NSCs have higher QDS modes than young NSCs, and the QDS modes decrease as cells gradually progress from dormancy to activation (similar to Fig. 4).

## 4 Notes

1. R is a programming language for statistical computing. RStudio is an integrated development environment for running R code. To download R and RStudio, go to https://posit.co/download/rstudio-desktop/; follow the installation steps.
2. To download the QDSWorkflow package, go to https://github.com/micw42/QDSWorkflow; follow the installation steps.
3. The data should be raw counts from RNA sequencing, with rows as genes and columns as samples. If the data is in plain text format (i.e., .txt, .csv file), load it into R using the data.table::fread() function. If the data is in the .rds format, load it into R using the readRDS() function.
4. The **BulkPreproc()** function preprocesses bulk RNA-seq raw counts. It normalizes and variance-stabilizes the raw counts using the varianceStabilizingTransformation() function from the DESeq2 package [24]. It then subsets the normalized counts to the genes that have higher than a threshold number of counts per million (CPM) in a threshold number of samples. Set the CPM threshold using the counts_thresh argument and the number of samples using the cell_thresh argument. As default, counts_thresh=0.5 and cell_thresh=2.
5. The **FilterDoublets()** function uses the DoubletFinder package [25] to find and remove doublets. Since the proportion of homotypic doublets (multiple cells of the same type sequenced together) may vary across scRNA-seq samples, the following steps run on each sample separately. I) To find the significant principal components (PCs), PCA is performed using the NormalizeData(), FindVariableFeatures(), ScaleData(), and RunPCA() functions from the Seurat package [26]. Typically, there are around 6-10 significant PCs. II) To find the optimal neighborhood size, a parameter sweep is performed on the significant PCs using the paramSweep_v3(), summarizeSweep(), and find.pK() functions from DoubletFinder. III) To cluster the cells, the significant PCs are used in the RunUMAP(), FindNeighbors(), and FindClusters() (with the "resolution" parameter set as 0.1) functions from Seurat. Typically, there are around 1-3 clusters. IV) The clusters are used in the modelHomotypic() function from DoubletFinder to estimate the homotypic doublet rate. V) The homotypic doublet rate and optimal neighborhood size are used to estimate the number of doublets. VI) The doubletFinder_v3() function from DoubletFinder is used to find the doublet cells, with the "PCs" parameter set to the number of significant PCs, the "pK" parameter set to the optimal neighborhood size, and the "nExp" parameter set to the estimated number of doublets. When running the FilterDoublets() function, set the following arguments: ann_df (a table with the sample label of each cell), split.by (the column in ann_df containing the sample labels), and id_col (the column in ann_df containing the cell IDs). Clustering fails on samples with 50 cells or less; the doublet-finding algorithm is not run on these samples, and a warning is given. FilterDoublets() can take several minutes or longer to run on datasets with more than 5000 cells.
6. Choose the threshold QC metrics by plotting the distributions of the total raw counts (nUMI), total number of non-zero genes (nGene), number of non-zero genes per total raw counts (log10GenesPerUMI), and mitochondrial gene counts per total raw counts (mitoRatio) using the **MakeHist()** function. Use these histograms to decide the threshold for each QC metric. Be relatively permissive; avoid filtering out cells that are part of continuous distributions but filter out only outlier peaks. For example, Fig. 6 shows the nGene distribution of dataset 2.2.2c. There is a small outlier peak at around 820, so we set nGene > 820 as the threshold.
7. The **scPreproc()** function preprocesses scRNA-sequencing raw counts. First, it filters out low-quality cells by QC threshold values (see note 6). Then, it normalizes the raw counts using the SCTransform package (with "vst.flavor" set as "v2") [27], which removes the effect of technical confounders using regularized negative binomial regression. Finally, it filters out low-expression and low-variability genes. To filter low-quality cells, set the QC thresholds using the nUMI_filt, nGene_filt, log10_filt, and mito_filt arguments. As default, nUMI_filt=500, nGene_filt=250, log10_filt=0.75, and mito_filt=0.2. Set the SCTransform variability threshold using the variable.features argument. As default, variable.features=10000, so SCTransform keeps the top 10000 most variable genes. To filter low-expression genes, set the threshold proportion of cells using the prop argument. As default, prop=0.002, so genes expressed in fewer than 0.2% of cells are filtered. This function can take several minutes to run on datasets with more than 5000 cells.
8. The training data should contain RNA-sequencing samples with at least 2 different dormancy depth levels. Bulk and scRNA sequencing data can both be used, but bulk is recommended because scRNA-sequencing data is often sparse and noisy. The sample labels should be numerical values corresponding to cell dormancy depth. The QDSWorkflow package comes with the training dataset 2.2.1; access it using the variable "ref_raw". Alternatively, an external dataset can be loaded into R (see note 3) and used for training.
9. Use the g:Profiler tool [28] to convert gene names to Ensembl ID. Go to https://biit.cs.ut.ee/gprofiler/convert. Copy the gene symbols/Ensembl IDs and paste them into the "Query" text box. Select the correct Organism, keep "Target namespace" as "ENSG" and "Numeric IDs treated as" as "ENTREZGENE_ACC." Click "Run query." When the query is finished, click "Export to CSV" to download the gene table as a csv file. Then, load the gene table into R (using the data.table::fread() function) and run **convertSymbol()**. Set the gene_map argument as the gene table, conv_col as the column with the converted IDs, and init_col as the column table with the original IDs.
10. Use the BioMart tool [29] to convert Ensembl ID between species. Go to http://www.ensembl.org/biomart/martview. Under "Choose Database," select "Ensembl Genes 109". Under "Choose Dataset," select the original species. Under "Filters," go to "Multiple Species Comparisons." Check "Homologue Filters" box. Use the drop-down menu to select the second species. Select "Only." Under "Attributes," select "Homologues (Max select 6 orthologues)." Find the orthologues of the second species. Check the "Gene stable ID," "orthology confidence [0 low, 1 high]," and "homology type" boxes. Click "Results", and check "Unique results only." Next to "Export results to," select "File" and "TSV." Click "Go" to download the homology table. Then, load the homology table in R (using the data.table::fread() function) and run **convertSpecies()**. Set the hm argument as the homology table, old_id as the column containing the original species IDs, and new_id as the column containing the new species IDs. For convenience, the QDSWorkflow package includes the rat-mouse and rat-human homology tables. Access them using the variables "rat_mouse_hm" and "rat_human_hm," respectively.
11. The **BootstrapSamples()** function first subgroups the cells in each sample by cell type; it then generates bootstrapped samples by sampling n cells with replacement from each subgroup for m times. Next, it bins each bootstrapped sample by taking the sum of the normalized counts per gene in the n cells. Set the ann_df argument as a table with the sample and cell type labels for each cell. For public datasets, cell-type labels are usually provided in the metadata, or available upon request to the author. For experimental data, cell-type labels can be inferred using Seurat clustering [26]. Set the group_col argument as the column(s) in ann_df containing the sample and cell type labels. Set the n argument as the number of cells in a bootstrap sample (as default, n=50). Set the m argument as the number of bootstrap samples to generate for each cell type in each scRNA-seq sample (as default, m=100).
12. The **QDS()** function merges the training and test datasets and removes batch effects between them using the ComBat() function from the sva package [22]. It sets the "batch" parameter as a vector indicating whether each sample is training or test. Then, it uses the cv.glmnet() function from the glmnet package [23] to optimize the alpha parameter (see note 13) and build an elastic net regression model on the training data. For the cv.glmnet() function, the "x" parameter is set as the batch-corrected training data, the "y" parameter is set as the training labels, the "type.measure" parameter is set as "mse," and the "nfolds" parameter is set as the number of training samples. Finally, it uses the model to predict on the test data, and outputs a table containing the ID of each test sample and its corresponding quiescence depth score (QDS). Provide the following arguments: test_df (the normalized test data), train_df (the normalized training data), ann_df (a table with labels of the training data), and y_col (the column in ann_df that contains the labels). Note that the predicted QDS values in the test dataset reflect cell dormancy depth mapped to the training model across different cell types and experimental conditions (and even different species). Therefore, the predicted QDS difference in the test dataset should be interpreted as the relative difference in cell dormancy depth between test samples.
13. In elastic net, the alpha parameter controls how many coefficients (genes) are in the model: lower alpha gives models with more coefficients, higher alpha gives models with fewer coefficients [23]. The **QDS()** function tests a range of alpha values and finds their associated cross-validation (CV) errors. To capture a broad gene signature of quiescence depth, we build the training model using the minimum alpha with a CV error within 5% of the minimum CV error.
14. The **make_boxplot()** function groups the test samples by condition and makes a boxplot of the QDS values in each group. Set the df argument as the table of predicted QDS values. Set the ann_df argument as a table with the condition label of each test sample. Set the x argument as the column in ann_df that contains the condition labels. Set y as the column in df that contains the QDS values. As default, y="QDS". Set the color argument as the column in ann_df to use when coloring the points (optional). Set the title argument as the title of the plot (optional). Set the levels argument as the order of the groups on the plot (optional).
15. The **make_grouped_boxplot()** function generates a boxplot of the QDS values in each subgroup of a given condition. Set the df argument as the table of predicted QDS values. Set the ann_df argument as a table with the condition and subgroup labels of each test sample. Set the x1 argument as the column in ann_df that contains the condition labels. Set the x2 argument as the column in ann_df that contains the subgroup labels. Set y as the column in df that contains the QDS values. As default, y="QDS". Set the title argument as the title of the plot (optional). Set the levels argument as the order of the groups on the plot (optional).
16. The **make_grouped_hist()** function plots a density curve for the predicted QDS values of each scRNA-seq cell population. Set the pred_df argument as the table of predicted QDS values. Set the ann_df argument as the table with the sample label of each cell. Set the grouping1 argument as the column in ann_df that contains the sample labels. Set the xvar argument as the column in pred_df containing the predicted QDS values. As default, xvar="QDS". Set the grouping2 argument as the column in ann_df that contains the subgroup labels (optional). Set the title argument as the title of the plot (optional). Set the levels argument as the order of the density curves on the plot (optional).

**Fig. 6.**
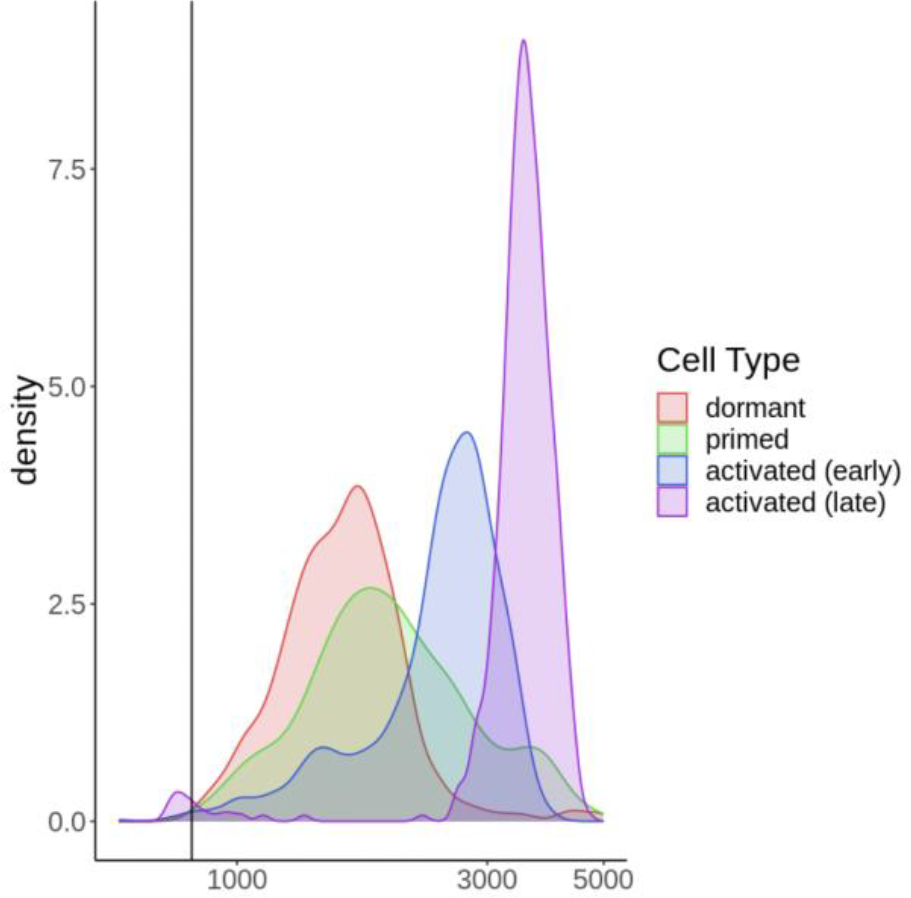
Histogram of the number of non-zero genes in the NSC dataset (2.2.2c) [21]. The vertical line represents the filtering threshold of nGene > 820. This threshold is chosen to filter out a small outlier subpopulation of the activated (late) cells (green distribution curve) while minimizing the removal of cells from other continuous cell distribution curves.

## References

1. Sender, R. and R. Milo, The distribution of cellular turnover in the human body. Nat Med, 2021. 27(1): p. 45–48.

2. Coller, H.A., L. Sang, and J.M. Roberts, A New Description of Cellular Quiescence. PLoS Biol, 2006. 4(3): p. e83.

3. Yao, G., Modelling mammalian cellular quiescence. Interface Focus, 2014. 4(3): p. 20130074.

4. Augenlicht, L.H. and R. Baserga, Changes in the G0 state of WI-38 fibroblasts at different times after confluence. Exp Cell Res, 1974. 89(2): p. 255–62.

5. Martin, R.G. and S. Stein, Resting state in normal and simian virus 40 transformed Chinese hamster lung cells. Proc Natl Acad Sci U S A, 1976. 73(5): p. 1655–9.

6. Owen, T.A., D.R. Soprano, and K.J. Soprano, Analysis of the growth factor requirements for stimulation of WI-38 cells after extended periods of density-dependent growth arrest. J Cell Physiol, 1989. 139(2): p. 424–31.

7. Yanez, I. and M. O’Farrell, Variation in the length of the lag phase following serum restimulation of mouse 3T3 cells. Cell Biol Int Rep, 1989. 13(5): p. 453–62.

8. Gunther, G.R., J.L. Wang, and G.M. Edelman, The kinetics of cellular commitment during stimulation of lymphocytes by lectins. J Cell Biol, 1974. 62(2): p. 366–77.

9. Bucher, N.L., Regeneration of Mammalian Liver. Int Rev Cytol, 1963. 15: p. 245–300.

10. Adelman, R.C., et al., Age-dependent regulation of mammalian DNA synthesis and cell proliferation In vivo. Mechanisms of Ageing and Development, 1972. 1: p. 49–59.

11. Rodgers, J.T., et al., mTORC1 controls the adaptive transition of quiescent stem cells from G0 to G(Alert). Nature, 2014. 510(7505): p. 393–6.

12. Llorens-Bobadilla, E., et al., Single-Cell Transcriptomics Reveals a Population of Dormant Neural Stem Cells that Become Activated upon Brain Injury. Cell Stem Cell, 2015. 17(3): p. 329–40.

13. Rodgers, J.T., et al., HGFA Is an Injury-Regulated Systemic Factor that Induces the Transition of Stem Cells into GAlert. Cell Rep, 2017. 19(3): p. 479–486.

14. Lee, G., et al., Fully reduced HMGB1 accelerates the regeneration of multiple tissues by transitioning stem cells to GAlert. Proc Natl Acad Sci U S A, 2018. 115(19): p. E4463–E4472.

15. Baker, D.J. and J.M. Sedivy, Probing the depths of cellular senescence. J Cell Biol, 2013. 202(1): p. 11–3.

16. Fujimaki, K., et al., Graded regulation of cellular quiescence depth between proliferation and senescence by a lysosomal dimmer switch. Proc Natl Acad Sci U S A, 2019. 116(45): p. 22624–22634.

17. Fujimaki, K. and G. Yao, Cell dormancy plasticity: quiescence deepens into senescence through a dimmer switch. Physiol Genomics, 2020. 52(11): p. 558–562.

18. Kwon, J.S., et al., Controlling Depth of Cellular Quiescence by an Rb-E2F Network Switch. Cell Rep, 2017. 20(13): p. 3223–3235.

19. Marthandan, S., et al., Conserved Senescence Associated Genes and Pathways in Primary Human Fibroblasts Detected by RNA-Seq. PLoS One, 2016. 11(5): p. e0154531.

20. Zhang, X., et al., The loss of heterochromatin is associated with multiscale three-dimensional genome reorganization and aberrant transcription during cellular senescence. Genome Res, 2021. 31(7): p. 1121–35.

21. Kalamakis, G., et al., Quiescence Modulates Stem Cell Maintenance and Regenerative Capacity in the Aging Brain. Cell, 2019. 176(6): p. 1407–1419.e14.

22. Leek, J.T., et al., The sva package for removing batch effects and other unwanted variation in high-throughput experiments. Bioinformatics, 2012. 28(6): p. 882–3.

23. Friedman, J., T. Hastie, and R. Tibshirani, Regularization Paths for Generalized Linear Models via Coordinate Descent. J Stat Softw, 2010. 33(1): p. 1–22.

24. Love, M.I., W. Huber, and S. Anders, Moderated estimation of fold change and dispersion for RNA-seq data with DESeq2. Genome Biol, 2014. 15(12): p. 550.

25. McGinnis, C.S., L.M. Murrow, and Z.J. Gartner, DoubletFinder: Doublet Detection in Single-Cell RNA Sequencing Data Using Artificial Nearest Neighbors. Cell Syst, 2019. 8(4): p. 329–337.e4.

26. Hao, Y., et al., Integrated analysis of multimodal single-cell data. Cell, 2021. 184(13): p. 3573–3587.e29.

27. Hafemeister, C. and R. Satija, Normalization and variance stabilization of single-cell RNA-seq data using regularized negative binomial regression. Genome Biol, 2019. 20(1): p. 296.

28. Reimand, J., et al., g:Profiler--a web-based toolset for functional profiling of gene lists from large-scale experiments. Nucleic Acids Res, 2007. 35(Web Server issue): p. W193–200.

29. Smedley, D., et al., BioMart--biological queries made easy. BMC Genomics, 2009. 10: p. 22.

